# A deep exon cryptic splice site promotes aberrant intron retention in a von Willebrand disease patient

**DOI:** 10.1101/2021.11.02.466821

**Authors:** John G. Conboy

**Affiliations:** Biological Systems and Engineering Division, Lawrence Berkeley National Laboratory, Berkeley, CA

## Abstract

A translationally silent single nucleotide mutation, in exon 44 of the von Willebrand factor (VWF) gene, is associated with inefficient removal of intron 44 in a von Willebrand disease (VWD) patient. This intron retention (IR) event was previously attributed to altered secondary structure that sequesters the normal splice donor site. We propose an alternative mechanism: that the mutation introduces a cryptic splice donor site that interferes with function of the annotated site to favor IR. We evaluated both models using minigene splicing reporters engineered to vary in secondary structure and/or cryptic splice site content. Analysis of reporter splicing efficiency in transfected K562 cells suggested that the mutation-generated internal splice site was sufficient to induce substantial IR. Mutations predicted to vary secondary structure at the annotated site had modest effects on IR, and also shifted the balance of residual splicing between the cryptic site and annotated site, supporting competition between the sites. Further studies demonstrated that introduction of cryptic splice donor motifs at other positions in E44 did not promote IR, indicating that interference with the annotated site is context-dependent. We conclude that mutant deep exon splice sites can interfere with proper splicing by inducing IR.

## Introduction

Aberrant splicing is a common cause of human genetic disease, with various studies estimating that 15-60% of disease-causing mutations disrupt splicing ([1, 2]; reviewed in [3]). Moreover, 5-20% of cancer-predisposing mutations can adversely affect splicing patterns [4]. Mechanistically, mutations can create new splice site motifs, alter enhancer or silencer elements that promote or inhibit recognition of splice sites, or alter the function of splicing regulatory proteins ([5–9] and references therein). Mutations can also alter RNA secondary structure to modulate accessibility of these motifs or binding affinity for relevant RNA binding proteins [10–15]. Aberrant effects on the transcriptome manifest as whole or partial exon skipping, exonification of intron sequences, and whole or partial intron retention. In some cases, even synonymous single nucleotide exonic mutations that don’t impact known regulatory motifs can adversely affect pre-mRNA splicing patterns. Distinguishing the mechanisms by which such mutations disrupt splicing requires detailed studies that ultimately will improve therapeutic strategies.

Mutations that impact RNA splicing at the level of intron retention have received relatively little attention. Recently, intron retention was reported in a patient with a bleeding disorder known as type I von Willebrand disease (VWD), caused by deficiency of von Willebrand factor (VWF) protein. A translationally silent C T mutation in exon 44 of the VWF gene in that patient, which substituted one glycine codon for another, is associated with altered splicing [16]. Although the mutation maps 86nt from the affected splice site, it nevertheless acts from a distance to cause retention of downstream intron 44 [16]. Computer modeling suggested that the mutant RNA adopts an aberrant secondary structure, sequestering the annotated 5’ splice site of exon 44 in a double-stranded configuration that renders intron 44 splicing inefficient [16]. Retention of the 2.2kb intron 44 sequence introduces premature termination codons into the aberrant transcript and would prevent synthesis of full length protein from the mutant allele, consistent with the observed VWF deficiency phenotype. However, no experimental testing of the secondary structure model has been reported.

The current study examines an alternative model for intron retention in this VWF patient, based on the observation that the mutation generates a strong 5’ splice site motif deep within exon 44. The presence of cryptic splice site motifs near annotated exon/intron boundaries has been shown in several contexts to inhibit use of neighboring annotated sites, leading to intron retention [17–19]. Here, using a minigene splicing reporter that reproduces VWF intron 44 retention, we engineered numerous mutations designed to distinguish whether altered secondary structure, or introduction of a new splice site motif, is the primary determinant of intron retention. Collectively the data suggest not only that secondary structure can impact intron retention, but also that the internal E44 splice site is a strong inducer of IR independent of secondary structure considerations.

## Results

VWF exon 44 normally possesses a fairly weak 5’ splice site, AGT/gtaggt, which has a score of 3.31 according to a commonly used algorithm for estimating splice site strength [20]. Inspection of exon 44 sequence revealed the presence of a very weak potential 5’ splice site motif (AAG/gcgagt; score=2.72), deep in the exon. This motif would likely be nonfunctional in normal individuals, and indeed there is no evidence for splicing at that site. However, the patient’s C→T mutation created a GT dinucleotide that greatly strengthens this motif to a near-consensus splice site (AAG/gtgagt; score=10.67). Weak 5’ splice sites are often susceptible to regulation generally, and splice sites flanking retained introns often reside near potentially competing splice sites [17–19]. We therefore hypothesized that the patient’s mutation induces intron retention not only by altering RNA secondary structure, but also via creation of a strong internal 5’ splice site.

To investigate the mechanism(s) responsible for IR in the VWF gene, we generated minigene splicing reporters whose splicing phenotype could be assessed following transfection into K562 erythroleukemia cells. SF3B1-mut35 is a splicing reporter used to study of intron 4 retention in the SF3B1 gene [17]. This construct is a variant modified to remove all of its decoy exons, features that promote IR for a subset of retained introns [17]. It can function as an IR reporter, i.e., retention is observed if heterologous IR-promoting elements are cloned into the intron.

Two series of VWF splicing reporters were engineered into the mut35 base construct. VWFwt-short was generated by inserting into mut35, at the site previously occupied by decoy exon 3e [17], a 314nt region of normal VWF genomic sequence spanning exon 44 and short flanking intron sequences (Figure 1A). A variant reporter containing the patient’s mutation, VWFm1-short, was altered at the indicated site. These reporters were designed to test whether intron retention might occur via the decoy exon mechanism, which would require only sequences proximal to E44. A second set of longer reporters was constructed to test whether IR only occurs in the context of full length intron 44 (Figure 1B). The long reporters contained 2.65kb of VWF genomic sequence extending from the distal portion of intron 43 to a proximal region of intron 45.

**Figure 1.**
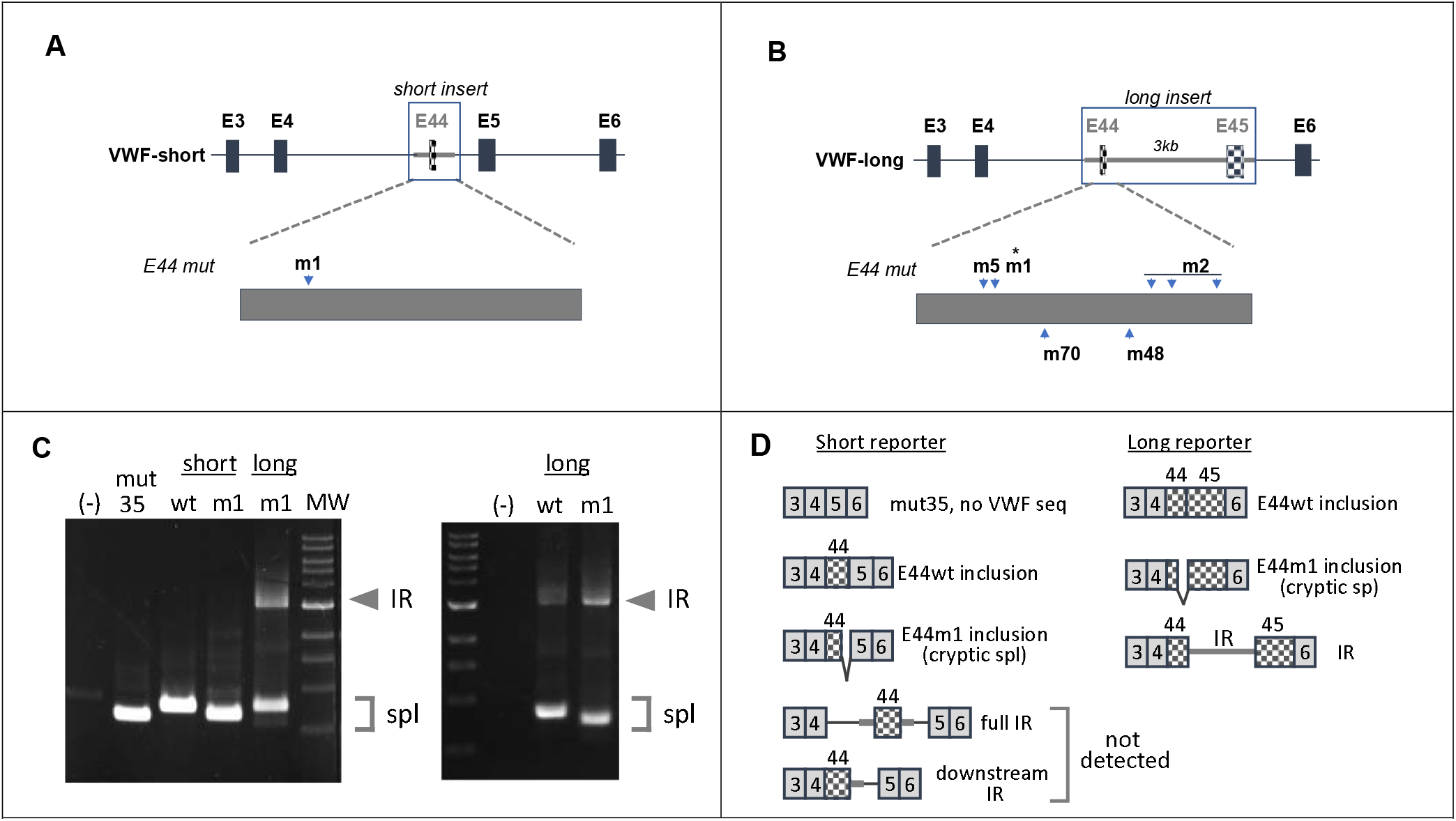
Engineering patient-specific retention of VWF intron 44 in minigene splicing reporters. A. Structure of short reporter. B. Structure of long reporter. C. Gel analysis of spliced products in K562 cells generated from parent construct lacking VWF sequences (mut35); short constructs containing normal (wt) or patient (m1) sequences for VWF E44 with short flanking intronic elements; and long constructs containing normal (wt) or patient sequences for including E44, intron 44, and E45. Position of additional mutations to be discussed later is indicated. D. Expected spliced products from short (left) and long (right) reporters. IR was primarily detected in transcripts derived from long reporters with the m1 mutation.

Gel analysis of spliced products amplified from transfected K562 cells is shown in Figure 1C, and the structures of these products deduced after sequencing are depicted in Figure 1D. As expected, the base construct mut35 did not exhibit IR. Similarly, neither of the VWF-short reporters, containing E44wt or E44m1 sequences, yielded any evidence for IR. A positive control containing the OGT decoy exon, processed in parallel and analyzed on a separate gel, did show substantial IR (data not shown). The absence of IR transcripts derived from these short VWF constructs suggests that E44 does not function as a decoy exon to promote IR. Regarding spliced products, VWFwt-short product was larger than for mut35 due to the inclusion of VWF E44, and the spliced m1 transcript was slightly smaller due to utilization of the cryptic m1 deep exon 5’ splice site. This result not only confirmed potential of m1 to recognized as a functional splice site, but also suggested that it out-competes the annotated site, perhaps due to the predicted secondary structure-based inaccessibility of the latter. Splicing at the m1 splice site was not reported in the patient, likely because the aberrant splice would have altered the translational reading frame to induce nonsense mediated decay.

Having reproduced the VWF patient’s IR phenotype, we then generated a series of splicing reporters to explore the mechanism(s) by which mutation m1 promotes IR. These reporters were designed to alter either of the features proposed to induce IR: (a) secondary structure at the annotated E44-5’ss that might impact its ability to interact with the splicing machinery [16], and/or (b) aberrant internal 5’ splice site motif(s) that might compete with the annotated 5’ss for interaction with that machinery (e.g., [19]). The predicted secondary structures for normal E44 and patient E44m1 are shown in Figure 2. These were adapted from [16], with the addition of stem designations to facilitate discussion of various structures. In normal E44, the weak internal 5’ss motif is located in the left arm of stem 1, with the annotated E44-5’ss predicted to be downstream of stem 2 in a relatively open (single-stranded) conformation (upper left). In the patient, reordered folding was previously predicted to incorporate the cryptic internal 5’ss, m1, into the right arm of new stem 3, with the annotated 5’ss partially sequestered in new stem 5 (upper right). Because the stabilities of these structures are not dramatically different [16], it seems plausible that the two might exist in some equilibrium within normal cells, and that E44 mutations could alter their relative abundance to impact IR.

**Figure 2.**
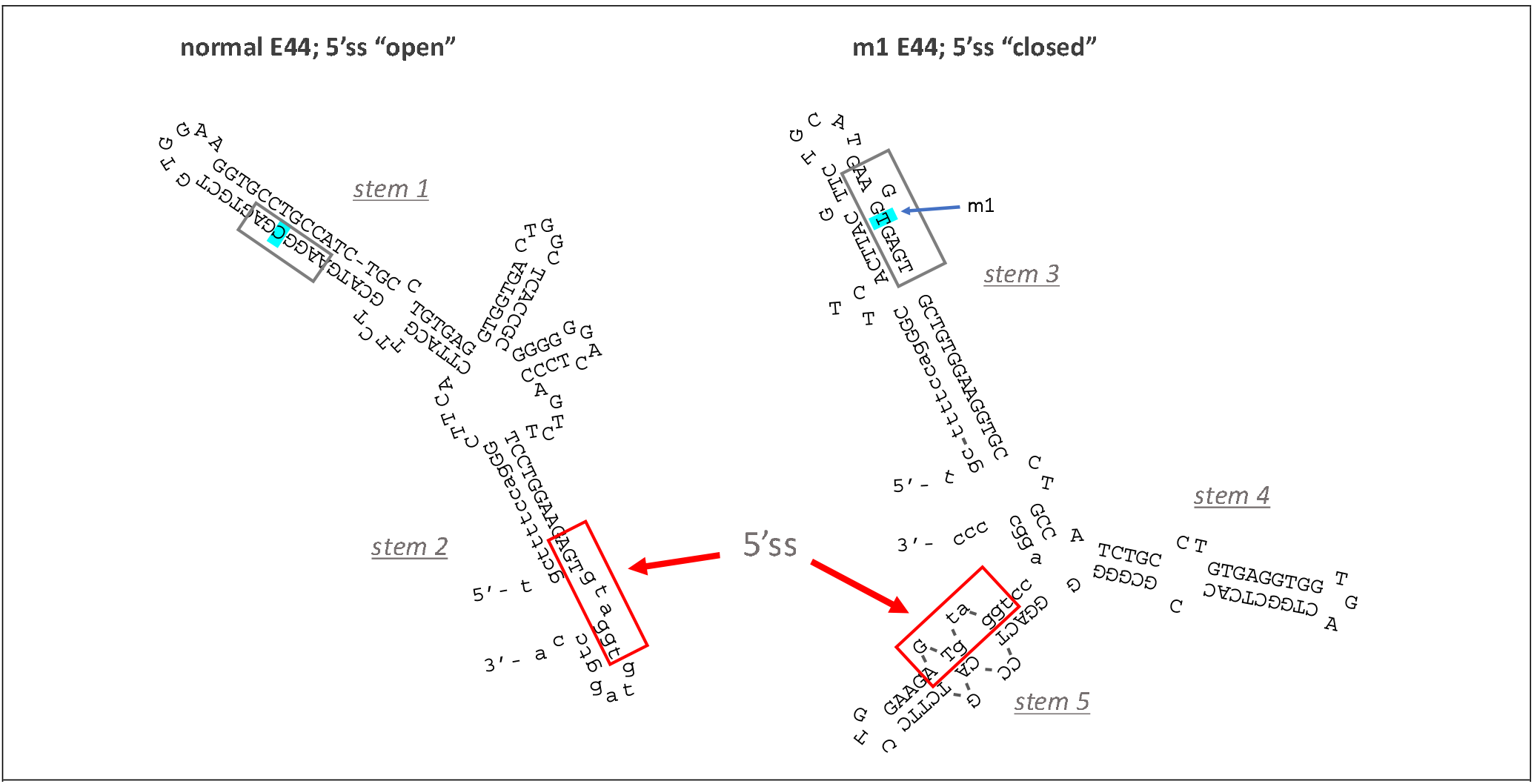
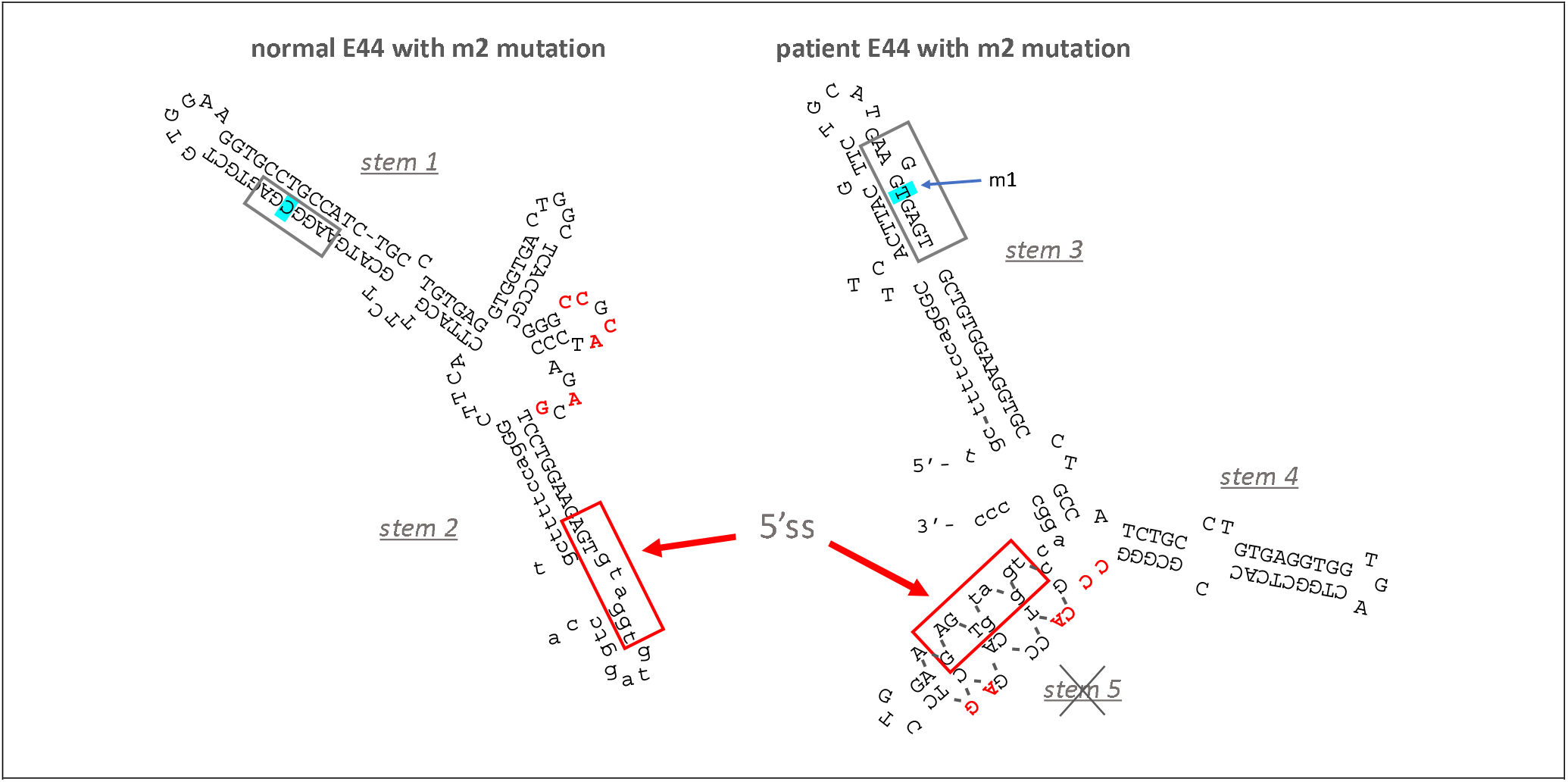
(A). Predicted structure of wild type E44 (“open 5’ss”) and patient E44 (“closed 5’ss”), redrawn from [16]. 5’ splice site motifs are boxed and annotation of stem structures has been added for clarity. (B). Superimposition of m2 mutations (red text) on normal “open” and patient “closed” conformations for E44 shown in Figure 2A. M2 mutations are predicted to disrupt stem 5, increasing accessibility of the 5’ss.

Reporter VWFm2 was designed to test the hypothesis that secondary structure at the annotated site is the major driver of intron 44 retention. VWFm2 contains six nucleotide substitutions expected to disrupt stem 5, thereby reducing the potential for secondary structure at the annotated 5’ splice site (Figure 2B, right). According to this model, an open conformation should greatly reduce IR, whether or not cryptic site m1 is present. Analysis of splicing patterns in K562 cells transfected with VWFm2 and relevant control reporters yielded two important findings (Figure 3). First, although IR in VWFwt was quite modest, the proportion of retention product relative to spliced product generated from VWFm2 was even less (compare lanes wt and m2). Second, considerable retention was observed in double mutant VWFm1m2. This result indicates that the cryptic site can induce retention even when the annotated site is predicted to be relatively accessible. Regarding accessibility of the annotated site, there is suggestive evidence that m2 favors a more open conformation, given the finding that the residual splicing observed to occur at the cryptic site for m1 (see Figure 1C) was switched back to the annotated site in m1m2. These results support the hypothesis that secondary structure at the annotated 5’ss of E44 does impact intron retention, but the presence of an internal cryptic 5’ss also strongly induces IR.

**Figure 3.**
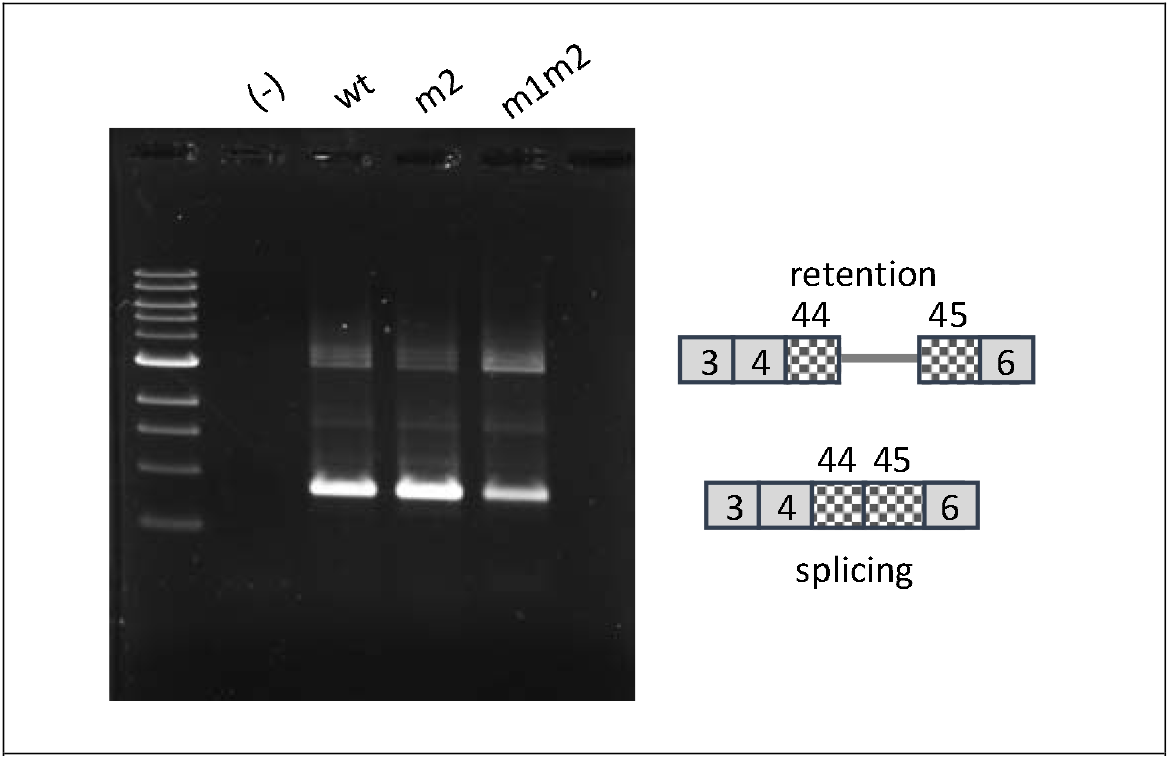
Effects of m2 mutations on IR. Lane (−), untransfected cells. Other lanes represent RT-PCR products from cells transfected with VWFwt, VWFm2, and VWFm1m2. Diagrams at right indicate structures of the spliced products.

Next, we generated a construct aimed at testing the hypothesis that inaccessibility of the annotated site alone can promote IR, even in the absence of the m1 cryptic site. Mutation m5 was predicted to mimic m1-mediated stabilization of stem 3, given the identical ΔG value of the two structures (Figure 4, left). M5 was therefore expected to favor the closed conformation associated with sequestration of E44-5’ss.

**Figure 4.**
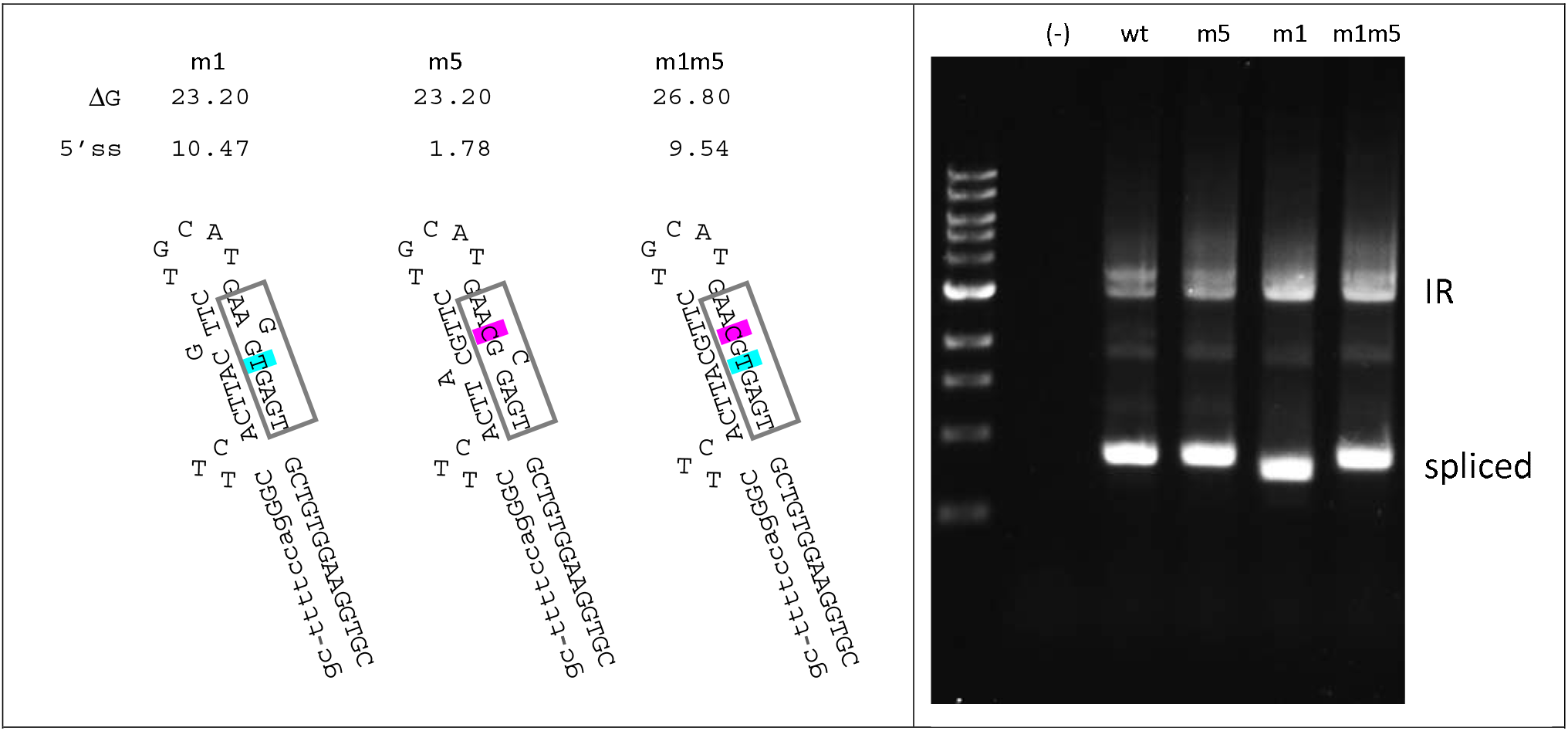
Effects of mutation m5 on IR predicted RNA structure and splice site strength (left) and splicing (right). For the gel analysis of splicing, lane (−) represents untransfected K562. Other lanes were transfected with VWFwt, VWFm5, VWFm1, and VWFm1m5 as indicated.

However, m5 differs from m1 in that it occurs in the context of a very weak 5’ splice site motif (AAC/gcgagt; score =1.78). Splicing analysis in K562 cells revealed that m5 yielded much less retention product relative to spliced product compared to m1, but slightly more retention product than VWFwt (Figure 4, right). This result is consistent with the interpretation that sequestration of E44-5’ss, in the absence of a competing 5’ splice site, increases intron retention only to a minor extent.

For completeness we also tested the double mutant m1m5, which has features similar to m1, i.e., a strong 5’ splice site motif (slightly weaker than m1), and strong stabilization of stem 3. When tested in K562 cells, m1m5 exhibited strong IR (Figure 4B, right), but it also switched back to use of the annotated 5’ splice site for residual spliced products. While this result is difficult to interpret in detail, since competition between splice sites is complex and may depend on poorly characterized accessory factors [21, 22], this finding may indicate that the m1m5 cryptic site competes less well with the annotated site either due to its slightly weaker splice site motif (Figure 4, left), to its potential for greater sequestration in a hyperstabilized stem 3, or both. Nevertheless, taken together these results suggest that both RNA folding and splice site competition contribute to intron 44 retention, but that splice site competition is likely the major determinant.

Finally, we asked whether the ability of a strong internal 5’ss motif to promote IR is position-dependent. We generated splicing reporters containing strong 5’ splice site motifs, identical to that of m1 (AAG/gtgagt), at two different locations within E44. One was 48nt upstream of the annotated E44-5’ss (m48) and the other was 70nt upstream (m70). The splicing phenotypes of these pre-mRNAs are shown in Figure 5. As before, construct VWFm1 exhibited much higher IR than did VWFwt. However, neither VWFm48 nor VWF70 displayed IR above the wild type background. That mutations m48 and m70 did generate functional 5’ splice sites was confirmed by sequence analysis showing that the E44 spliced products in these mutants primarily utilized the internal cryptic sites. We interpret these results to indicate that an internal 5’ss motif must reside in an appropriate exonic context in order to interfere with splicing at the annotated site to yield intron retention.

**Figure 5.**
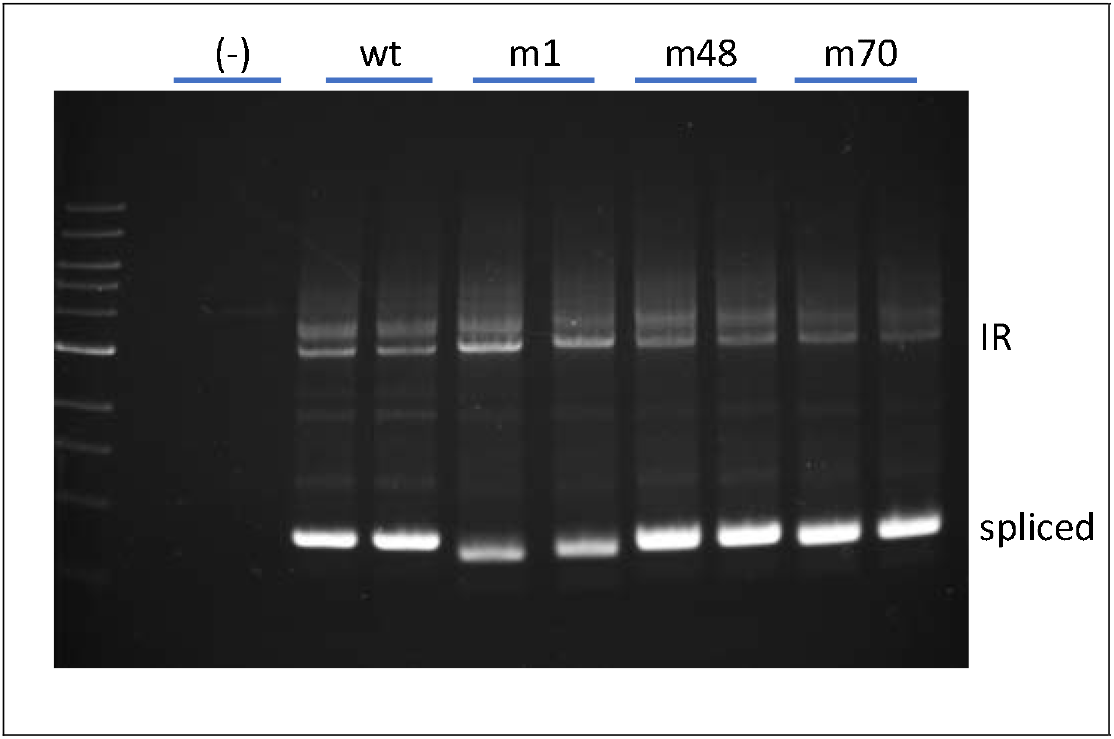
Position-dependence of IR stimulation by cryptic 5’ splice sites in E44. Analysis of splicing phenotype for E44 variants. Lanes (−) represent untransfected K562. Other lanes were transfected with VWFwt, VWFm1, VWFm48, and VWFm70.

## Discussion

Von Willebrand disease is one of the most common inherited bleeding disorders, caused by any one of hundreds of mutations in the VWF gene [23, 24]. The mutation studied here, although present at low frequency among the populations surveyed (dbSNP: <u>rs900907976</u>; minor allele frequency=0.000026 (7/264690, TOPMED), has an unusual impact on pre-mRNA splicing by acting at a distance to promote retention of the downstream intron [16]. Understanding how a “deep exon” mutation can promote IR may be important for design of therapies for patients with this and mechanistically-related diseases yet to be discovered.

Two major hypotheses have been proposed to explain how the m1 mutation interferes with function of the annotated 5’ splice site. First, the splice site sequestration model predicted that mutation m1 would alter long range RNA folding across exon 44 so as to increase base-pairing at the annotated 5’ splice site. According to this model, RNA-RNA interactions promote IR by sequestering that splice site away from the splicing machinery [16]. Second, the splice site competition model posits that function of the annotated site is compromised by the presence of the cryptic 5’ splice site created by m1. Productive splicing in this model would be silenced by protein-protein interactions between complexes bound at the decoy site within the exon and at the annotated site at the normal exon/intron boundary.

To distinguish between these models, we analyzed the behavior of splicing reporters altered at either of these features. One instructive finding was the difference in splicing behavior between mutants m1 and m5. Both were predicted to stabilize stem 3 (Fig. 4), favoring the conformation that sequesters the annotated 5’ splice site, but they differ greatly with regard to strength of the internal splice site motif. Mutant m1 possesses a strong splice site motif and exhibits high IR. In contrast, m5 has a very weak motif but equivalent sequestration potential, and it displays much lower IR. This finding indicates that sequestration alone is insufficient to induce substantial IR. A second critical comparison involved constructs m1 and m1m2. These have identical, strong, deep exon splice site motifs as described in the patient, but different sequestration potentials at the annotated site. Both exhibited strong IR, supporting the interpretation that the cryptic site is sufficient to induce IR. Taken together, these results indicate that secondary structure does impact splice site choice and IR; however, IR occurs even with a more open configuration at the annotated site, as long as an upstream cryptic site is present in an appropriate sequence context.

A potential limitation on these interpretations is that secondary structure has not been probed directly. However, using splice site usage as a proxy for accessibility, we observed that residual splicing switched from the cryptic site in m1 to the annotated site in m1m2, consistent with a more accessible conformation of the annotated site in the double mutant. Interestingly, the double mutant m1m5 shifts residual splicing in the opposite direction back to the annotated 5’ splice site. Although folding models suggest that m1 and m1m5 have equivalent potential to sequester the annotated site, the strong cryptic site in m1m5 may be disfavored for residual splicing either because it is slightly weaker than m1, or because m1m5 becomes sequestered by increased stability of stem 3 (Figure 4). A caveat to these interpretations is that secondary structure has a complex role in alternative splicing [13], and the predicted stability of stem loop structures may not be as critical as specific structures in determining RNA functionality [25].

Another limitation to this study is that we have not directly measured spliceosomal binding to the cryptic site. For that reason, the possibility that a non-spliceosomal complex may bind at the cryptic site region to induce IR cannot be formally excluded. However, several observations support involvement of a spliceosomal component. First, cryptic site m1 does operate as a functional splice site, as shown by sequence analysis of residual spliced products. Second, variant substrates bearing a strong cryptic splice site were associated with strong IR (m1, m1m2, m1m5), while those having a weak cryptic site were correlated with low IR (wt, m5). Finally, as discussed below, there is precedence for the idea that cryptic splice sites can interfere with annotated splice sites to promote IR [19].

In the Drosophila P element gene, binding of spliceosomal components to an exonic decoy splice site modulates retention of the downstream intron in a tissue-specific manner [19]. The splice site itself is necessary but not sufficient for regulation, with auxiliary RNA binding proteins being required to promote IR [26–28]. Binding of auxiliary protein factors can, in the presence of an exonic cryptic splice site, also inhibit productive splicing at the downstream authentic 5’ splice site to alter the balance of splice site choice [21, 29, 30]. In decoy exon-mediated intron retention, the decoy splice sites are often close to alternative splice site motifs that may play a role in enforcing nonproductive interactions with intron-terminal splice sites to inhibit intron excision [17, 18]. Even more distally, deep intron splice site motifs, defined via their binding to U2AF65, appear capable of competing with recognition of splice sites of neighboring downstream exons [31].

Given these precedents, we propose that the splice site introduced by the VWF patient’s mutation interacts in a nonproductive manner with the annotated 5’ splice site located 86nt downstream, likely via spliceosomal associations assisted by unknown accessory protein(s). If the downstream intron 44 is recognized by an intron definition mechanism [32], then blocking its 5’ splice site could yield IR. Not well understood yet is what neighboring sequence features constitute a permissive environment for induction of IR. Regardless, it’s interesting to speculate that blocking the cryptic splice site with antisense oligonucleotides might reactivate the annotated 5’ splice site to ameliorate the VWF protein deficiency.

More globally, relatively little is known about the incidence of IR due to deep exon mutations, and what might be their contribution to human disease. Methodological and conceptual considerations may have limited the identification of such mutations in previous exome and RNA-seq studies. Detecting association between deep exon SNVs and intron retention events requires reads of sufficient length to overlap both features, sufficient sequencing depth to identify allele-specific association of such features, and analytical strategies focussed on finding these associations. Earlier work has in fact revealed that many exonic SNVs are associated with IR [33], but this analysis was limited to exon sequences near (within 30nt of) exon-intron junctions. IR-associated exonic SNVs in that study mapped predominantly to the last nucleotide of the exon, and these likely act via direct interference with 5’ splice site recognition. A few IR-associated mutations were found outside of the splice site motif, but their mechanism of action was not explored other than to determine that they were not enriched in known splicing silencer or enhancer motifs [33]. Other studies have reported numerous deep exon mutations that create new 5’ splice site motifs and are associated with aberrant splicing [34]. These were detected in short RNA-seq reads that contain mutations linked to abnormal exon/exon junctions; whether any of these were also associated with IR was not assessed. Systematic analysis of deep exon mutations associated with IR might be enabled by acquisition of more long RNA-seq reads that overlap deep exon sequences and exon/intron boundaries. All of these analyses would likely benefit from application of algorithms that optimize detection of retained introns from short RNA-seq data [35, 36] and long RNA-seq data [37]. Finally, mutations that create novel 5’ splice site motifs and are associated with IR can also occur in downstream intron sequences [38].

## Materials and Methods

### Construction of splicing reporters

Reporters diagrammed in Figure 1 were constructed as follows by modification of an SF3B1-based minigene, mut35, in which intron 4 retention was suppressed due to removal of all decoy exons [17]. The first generation reporter (short) was made by amplifying a 314 nt region of VWF gene spanning exon 44 and proximal intron sequences using the following primers:

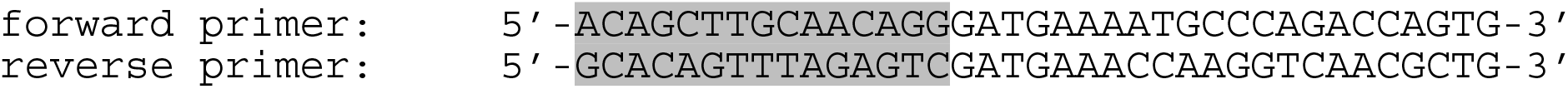

The shaded regions provided overlap sequences to facilitate fusion [39] of this fragment into mut35 at the normal position of decoy exon 3e [17].

The second wild type reporter (long) was made by using the following primers to amplify a 2.65kb region of the VWF gene spanning across a terminal portion of intron 43, exon 44, intron 44, exon 45, and a proximal region of intron 45.

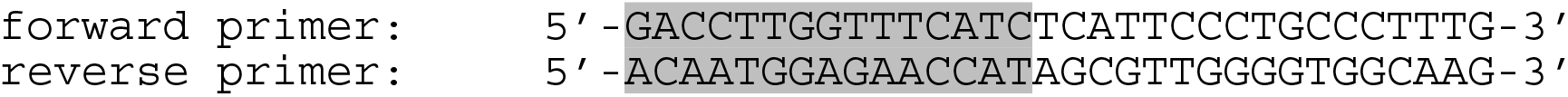

This fragment was cloned into mut35 using fusion methods [39] with 15 nt in lower case sequence representing overlap with the ends of the linearized mut35 vector. To accommodate this larger insert, mut35 was linearized by amplification using primers that omitted ~1.5kb of SF3B1 sequence (spanning the distal 0.2kb region of intron 4 to the proximal 1.3kb region of intron 5).

Mutations in VWF exon 44 were introduced by amplifying the wild type VWF reporter with primers carrying the desired mutations. The various 5’ splice site motifs (boxed) and their immediate flanking sequences are shown below. Construct m1 displays the patient’s sequence. Construct m5 was mutated to stabilize stem 3 equivalently to m1, but without generating a strong 5’ splice site motif (see Figure 4). m1m5 contains both the m1 and m5 mutations. Constructs m48 and m70 represent mutations in the wild type construct that introduce strong 5’ splice site motifs identical to that of m1. Asterisks indicate the G residues at the putative cryptic exon/intron boundaries. Shaded nucleotides differ from wild type sequence. The m1 splice site from the VWF patient is 86nt upstream from the annotated site, while m48 and m70 are 48nt and 70nt, respectively, upstream of the annotated site.

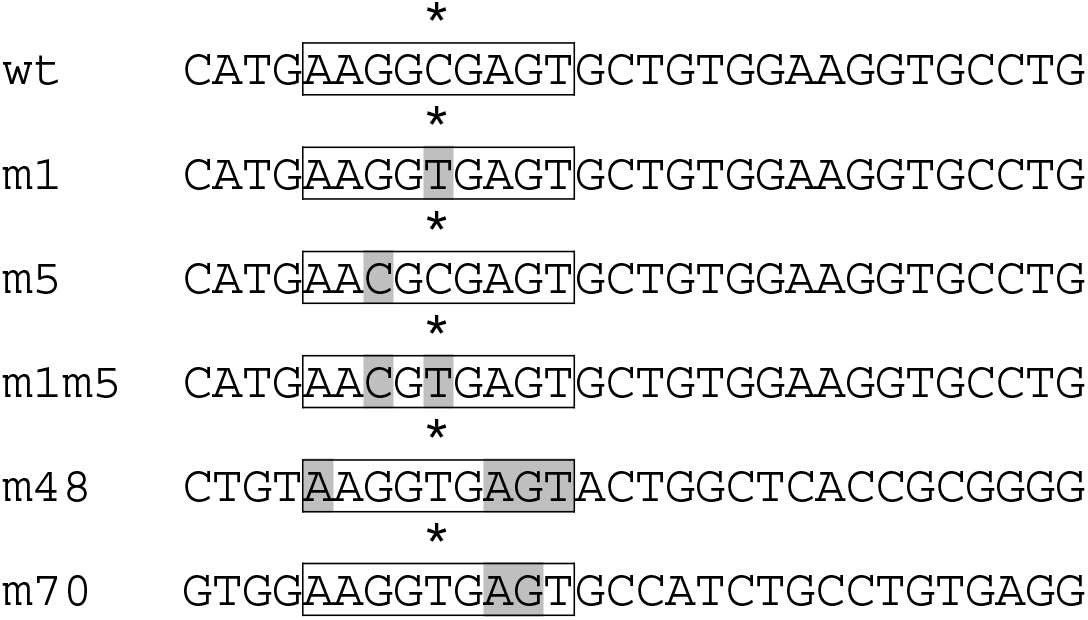

Construct m2 contains 6 nucleotide changes near the annotated 5’ splice site, designed to disrupt secondary structure in this region. The linear sequence of the annotated 5’ splice site region for the wild type (‘closed conformation’) and m2 (‘open conformation’) constructs is given below. Predicted secondary structures are shown in Figure 2.

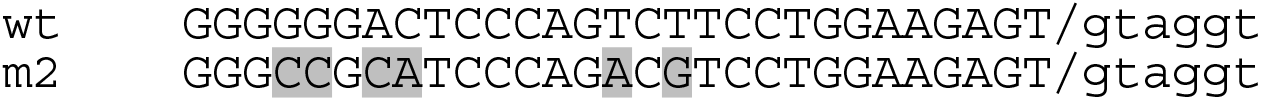

### Analysis of splicing

Splicing reporters were transfected into K562 using FuGENE according to the manufacturer’s instructions (Promega). RNA was isolated after 48 hrs using RNeasy columns as per instructions from the manufacturer (Qiagen), with the inclusion of a DNase step to minimize contamination by genomic DNA. RNA was reverse transcribed with Superscript III (Invitrogen) into cDNA using the BGH reverse primer in the vector (5’-tagaaggcacagtcgagg-3’). Spliced products were amplified using a forward primer in exon 3 (5’-catcatctacgagtttgcttgg-3’) and a reverse primer in the vector (5’-atttaggtgacactatagaatagggc-3’). This strategy amplified minigene-derived transcripts but not endogenous mRNA, as confirmed using RNA from untransfected or empty vector-transfected cells. When assaying IR products, PCR reaction conditions were adjusted to allow for amplification of DNA bands ≥3 kb in length (denaturation at 95°C for 20 sec, annealing at 56°C for 10 sec, extension at 70°C for 2 min 30 sec; 35 cycles) using KOD polymerase in the presence of betaine. PCR products were analyzed on either 2% agarose gels or 4.5% acrylamide gels. Identity of PCR products was confirmed by DNA sequencing, and all splicing reporters were assayed at least three times. The splicing phenotype of test constructs relative to control constructs was highly reproducible, despite variations in baseline intron retention.

## Acknowledgements

The author thanks UC Berkeley Cell Culture Facility, especially Alison N. Killilea and Molly Fischer, for considerable assistance with K562 cultures used in this study.

